# An *in vitro* neuronal model replicating the *in vivo* maturation and heterogeneity of perineuronal nets

**DOI:** 10.1101/2022.01.22.477344

**Authors:** S Dickens, A Goodenough, JCF Kwok

## Abstract

The perineuronal net (PNN) is a condensed form of extracellular matrix (ECM) that enwraps specific populations of neurons and regulates plasticity. To create a PNN, only three classes of components are needed: membrane bound hyaluronan by its synthetic enzyme hyaluronan synthases (HASs), a link protein and a CSPG. However, there is redundancy within the classes as multiple HAS isoforms, link proteins and CSPGs have been found in the PNN *in vivo*. The effect of this heterogeneity has on PNN function is unresolved. Currently, the most common way to address this question is through the creation and study of PNN component in knockout animals. Here, we reported the development of a primary neuronal culture model which reproduces the *in vivo* maturation and heterogeneity of PNNs. This model accurately replicated mature cortical PNNs, both in terms of the heterogeneity in PNN composition and its maturation. PNNs transitioned from an immature punctate morphology to the reticular morphology as observed in the mature CNS. We also observed a small population of PNNs that were mature at an earlier time point and a distinct composition, highlighting further heterogeneity. This model will provide a valuable tool for the study of PNN biology, their roles in diseases and the development of PNN focused plasticity treatment.

## 1.0 Introduction

The perineuronal net (PNN) is a key regulator of plasticity and conserved across species (*1–4*). It stabilises existing synapses at the end of critical period during development and restricts new synapses onto the enwrapped neurons (*5, 6*). Prolonging the critical period via monocular deprivation delays PNN development (*7*). PNN attenuation, through component knock out, prevents the closure of critical period (*8*). The low plasticity level in adult central nervous system depends on the PNN as degradation reawakens experience-dependent plasticity (*9*). Reawakening latent plasticity in the central nervous system (CNS) has great utility in disease treatment, e.g. spinal cord injury (*10*). Reawakening juvenile plasticity through PNN manipulation holds great hopes for treating spinal cord injury and stroke. Currently the only way to manipulate the PNN, without the use of transgenic technology, is through the generalised destruction of the ECM by enzymatic degradation (*11*). This can lead to maladaptive plasticity (*12*). Improving understanding of PNN development and how it exerts its function is key to a fine manipulation of plasticity.

The PNN is an extracellular structure and components originate from both the host neuron and other sources (*5, 13–15*). The surface anchors for the PNN must, by necessity, come from the neuron. HA chains are directly extruded from HA synthases (HAS) expressed by the PNN neuron (*13, 14, 16*). Until recently, HAS was the only known cellular anchor for the PNN (*17*). RPTPζ has been recently identified as a PNN anchor for aggrecan (ACAN) (*18*). Chondroitin sulfate proteoglycans (CSPGs) from the lectican family then bind to the hyaluronan (HA) chains. This interaction is stabilised by the hyaluronan and proteoglycan link proteins (HAPLNs). Trimeric glycoprotein tenascin-R (TnR) then binds to the C-terminal of the CSPGs, allowing crosslinking of CSPGs to take place. Both HAPLNs and TnR are expressed by the PNN-bearing neurons (*5, 13, 19*). The origin of CSPGs, on the other hand, depends on the individual CSPG. While both neurocan (NCAN) and ACAN originate from PNN-neurons, brevican (BCAN) is produced by both astrocytes and neurons (*5, 13, 20*). In addition to these core components, other ECM molecules could also bind and modulate PNN structure and functions. Semaphorin 3a (sema3A), Otx-2 and neuronal activity-regulated pentraxin (Narp) have been shown to bind to PNNs (*21–23*).

In addition to molecular heterogeneity, there is also a temporal difference in PNN development. The precise timing of PNN development varies between brain regions (*24, 25*) ranges from postnatal day (P) 7 and 56 in rodents and between 1 month to 20 years of age in humans. The development of PNN coincides with the closure of critical period (*13, 26*). As the PNN matures it transitions from a punctate morphology to a contiguous, reticular structure (*27, 28*). The punctate morphology represents an immature PNN phenotype as it is linked to persistent juvenile plasticity (*8, 25, 29*). During PNN development, the expression of PNN components rises and peaks. Peak timing varies between component and region (*14, 16, 30*). For example, in the cerebellum, *Ncan* mRNA expression peaks between P3-7 while *Acan* mRNA peaks later between P14-21 (*16*). The peak of a component even differs between brain areas; the *Ncan* peak occurs later in the cortex (P10) and spinal cord (P14-P21) (*14, 31*). The peak timing corresponds with PNN formation but a direct relationship between a single PNN component and PNN maturation has not been identified.

An *in vitro* neuronal PNN model would provide a platform to investigate the role of individual molecules on the functions and structures of PNNs. There are several advantages to PNN culture model: cost, time, ease of manipulation through genetic manipulation. While the effect on behaviour cannot be investigated, a neuronal PNN model would allow the molecular mechanisms to be determined with high resolution and precision. Previous *in vitro* PNN cultures have established the contribution of glia and neuronal activity to PNN formation (*5, 15, 32*). Some of the cultures have exhibited reticular morphologies but many have presented with punctate PNNs. In order to understand the structure-function relationship of PNN molecules, it is crucial to develop an *in vitro* PNN neuronal model which replicate the PNN heterogeneity and maturation.

In this project, we have demonstrated that *in vitro* maturation of E18 neuronal culture, supplemented with astrocyte condition medium, will allow for the development of PNN neurones, replicating the maturation, molecular heterogeneity and morphology of PNNs. Using this protocol, 86% of the PNN-positive neurons are with mature reticular PNN structure. These neurons demonstrate expression of ACAN, NCAN, BCAN, TnR, and HAPLN1, which corresponds to the endogenous development of PNNs. The model will be valuable for the study of PNN development and potential treatment for plasticity.

## 2.0 Methods

### 2.1 Neuronal culture

All animals were housed in standard housing conditions with a 12 hour light / dark cycle with food *ad libitum*. The time mated pregnant females were singly housed. The work was performed under the regulations of the Animals Scientific Procedures Act 1986 and covered by the Home Office project license (#70/8085) and PIL for Stuart Dickens (#I9C757118).

Cortical neurons from E18 Wistar rats. The pregnant female was humanely sacrificed according to the authorised scheduled 1 protocol in the UK before the embryo collection. The embryos were similarly sacrificed, the cortices were dissected out and suspended in cold Hank’s buffered salt solution, without ions (HBSS-) (ThermoFisher #11590466). The media was then aspirated and the cortices digested with filter-sterilised papain (2 mg/ml in HBSS-, Lorne Laboratories #LS003119) for 6 min at 37 °C. To degrade released DNA, DNase I was added to a final concentration of 50 μg/ml (Sigma #DN25). The cell containing supernatant was then taken and centrifuged at 100 xg, 2 min at room temperature. The cells were resuspended in Neurobasal™ media with 2%(v/v) B27, 0.4 mM Glutamax and 0.1% antibiotic-antimycotic (ThermoFisher #21103049, #17504044, #35050038, #15240062). Cells were plated at 100,000 cells/well in a 24 well plate containing 40 μg/mL poly-D-lysine treated coverslips.

After plating, half the neuronal media volume was replaced twice a week. For the first 7 days the neurons were fed with Neurobasal™ with additives. From 7-14 day *in vitro* (DIV), the media was replaced with BrainPhys™ medium with 2% (v/v) SM1 and antibiotic-antimycotic (StemCell Technologies #5792). From 14 DIV, the media was replaced with 1:1 BrainPhys™ media with SM1 and astrocyte conditioned BrainPhys™ media.

### 2.2 Astrocyte culture for conditioned medium

Astrocytes from P0-6 Wistar rats were harvested in accordance with the UK Home Office guidelines. The pups were humanely sacrificed as before. The cortices were then removed and manually dissociated with a sterile blade. The mixture was then digested for 20 min with a 0.1% filter-sterilised trypsin solution (Sigma #T0303) at 37 °C. DNase I was added as before. The cell containing solution was then centrifuged at 100 xg, 5 min at room temperature. The pellet was then resuspended in DMEM supplemented with 10% (v/v) foetal bovine serum (FBS) with antibiotic-antimycotic (Lonza #12-707F, ThermoFisher #15240062). The cells were then titrated with fire polished glass pipettes with different apertures. The cells were then plated in 40 μg/mL poly-l-lysine coated culture flasks. The media was then replaced the following day and every three days thereafter. Once the flasks were confluent, the cell population was enriched for astrocytes using differential adhesion. The flasks were shaken at 125 rpm overnight to remove contaminating cells. The astrocytes were then split and upscaled into larger flasks. Once the cultures reached 70-100% confluence, the media was replaced with BrainPhys with additives for 48 hrs. The conditioned media was then centrifuged to remove contaminating cells and filter sterilised.

### 2.3 Immunocytochemistry of neurones

Neurons were fixed at room temperature in 4% (w/v) paraformaldehyde, 3% (w/v) sucrose solution for 10 minutes. The cells were then washed in 3% (w/v) sucrose, permeabilised with 0.01%(v/v) Triton x-100 for 10 minutes. Non-specific binding sites were blocked by incubation with 3% (v/v) normal donkey serum (Sigma #D9663) in 1X Tris buffered saline (TBS) for 1 hr at room temperature. The neurons were then incubated with primary antibodies (see table 2), diluted in blocking solution, overnight at 4 °C. The following day the neurons were then washed with 1X TBS and incubated with the appropriate secondary antibodies, diluted in blocking solution, for 2 hours at room temperature. They were then washed in TBS and mounted onto SuperFrost microscope slides (Fisher Scientific Ltd #10149870) using FluoroSave (Merck #345789).

### 2.4 Imaging, quantification, and statistics

Imaging was performed either using a LSM880 confocal microscope (Zeiss) with 20x and 63x objectives or an AxioScan Z.1 Slidescanner (Zeiss) with a 20x objective. Coverslips were imaged with 20x objective and tile-scanning for quantification. High resolution images were created by z-stack imaging with 63x objective. A projection image was then created using max intensity plugin on ImageJ. For the Slidescanner, tile images with 20x objective were taken and stitched to provide entire coverslip image. Filters were set to ensure no overlap between channels. PNNs were identified and counted based on morphology. OriginPro 2019b was used in graph creation and statistical analysis. Tests for significance (p<0.05) were performed using one-way analysis of variance (*ANOVA*) with Tukey’s *post hoc* test.

## 3.0 Results

### 3.1 In vitro culture replicates PNN development in vivo

The development of the PNN is closely tied to the closure of the critical period and the maturation of PV neurons. Recent research has shown that critical period closure is not just dependent on the presence of PNNs but also of the morphology of the PNNs (*27, 28*). The contiguity and intensity of PNNs increases as they mature (Sigal, *et al.*, 2019). Previous *in vitro* PNN cultures have failed to replicate the reticular morphology seen in mature PNNs *in vivo*, only showing a punctate morphology (*22, 33, 34*). We hypothesised that the punctate PNNs is a temporary structure representing immature PNNs and a reticular morphology resembling adult PNN *in vivo* would develop over time (*35*).

We cultured primary rat cortical neurons until DIV56 and stained them for PNN components and morphology. The cultures were initially plated in Neurobasal based media to aid cell survival. At DIV7, the media was changed to BrainPhys based media to support neuronal activity (*36*). At DIV14, the media was again changed to astrocyte conditioned BrainPhys based media as astrocyte media is necessary for the long-term maintenance of cultures and astrocytes secrete several PNN molecules (Fig. 1A). Assessment of PNN formation was performed with WFA and ACAN staining as both are recognised markers of cortical PNNs. At DIV14, a small proportion of cortical neurons started to show weak punctate WFA staining. This became more widespread between DIV21 and DIV24 (Fig. 1B) which was then chosen as the earliest time point. PNNs initially presented with a punctate morphology, mimicking the immature pre-critical period PNN morphology seen *in vivo* (Fig. 1C, top). The PNN population transitioned to a reticular phenotype as the culture developed, with an intermediate phenotype seen between DIV28 and DIV35. (Fig. 1B). This PNN maturation mirrored the transition seen *in vivo* (Fig. 1C, P30). By DIV56, the PNNs had matured, with the majority presenting a reticular morphology (86%), mirroring the post-critical period mature PNNs at P90 *in vivo* (Fig. 1, C and D). However, as *in vivo,* a small proportion of PNNs remained punctate (Fig. 1D) (Sigal, *et al.*, 2019). Quantification of punctate and reticular PNN morphologies confirmed the maturation trend as shown by analysis of both WFA and ACAN positive PNNs (Fig. 1, D and E). The PNN population switched from a majority punctate to a majority reticular morphology between DIV28 and DIV35 the PNNs. This switch is reminiscent of the critical period closure observed *in vivo* indicating we have replicated PNN development *in vitro*. At DIV56, PNN neurons accounted for 7.62 ± 1.12% of total neurons. This is a similar proportion to the cortical PNN population *in vivo*, where PNN neurons account for 4-10% of total (*37, 38*). The majority of PNN neurons were reactive for parvalbumin (PV) (62.5 ± 0.075%), which compares to the cortical PNN population *in vivo* (*39*). Together this indicates that our culture mimics the cortical PNN population observed *in vivo*.

**Figure 1.**
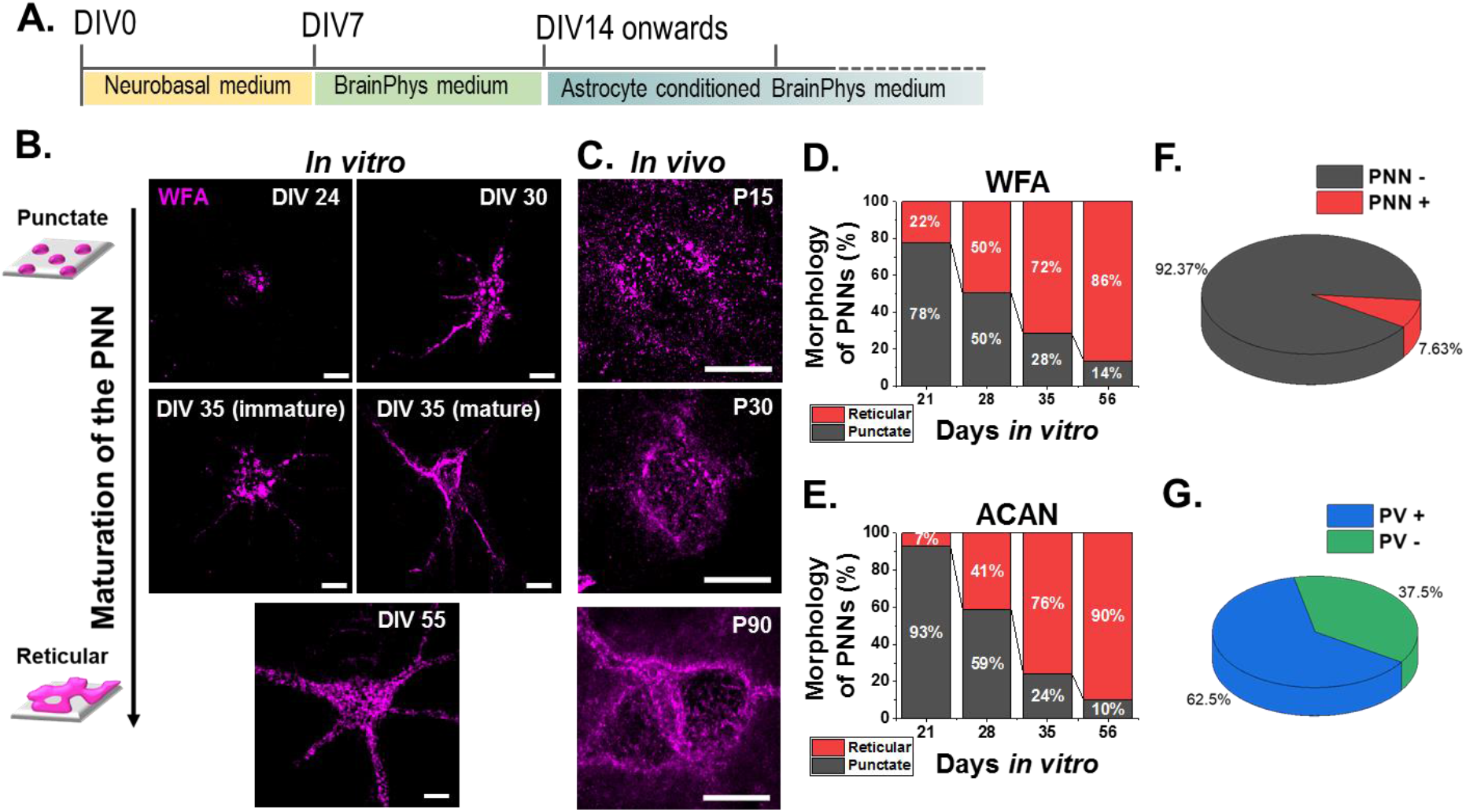
**(A)** A timeline showing the use of different culture medium for the long-term culture of cortical neurons for PNNs. **B)** Representative images of cortical PNN development *in vitro* as shown by WFA staining. **(C)** Representative images of *in vivo* PNNs at P15, 30 and 90. **(D)** Quantification of PNN morphologies as shown by WFA staining. **(E)** Quantification of PNN morphologies as shown by ACAN staining. Data from (D) and (E) are average percentages from 6 coverslips from 2 rats. **(F)** At DIV56, 7.63% ± 1.12% of neurons showed PNNs on surface. **(G)** In the PNN-positive neuronal population, 62.5 ± 0.075% of them are parvalbumin (PV) neurons. n= 3 rats. Scale bars in B and C are 10 μm.

### 3.2 PNN composition changes during development

Next, we investigated the changes of key PNN molecules during neuronal maturation. WFA is widely used as a pan-PNN marker and yet recent reports have shown that it does not identify all PNNs in the CNS (*40*). ACAN has been identified as another pan-PNN marker as it identifies a higher proportion of PNNs in both the cortex and the spinal cord (*40, 41*).

WFA staining recognised a majority of PNNs at all-time points despite their morphologies (WFA-positive neurons: DIV21: 71.3 ± 9.76%, DIV28: 58.4 ± 9.38%, DIV35: 71.2 ± 13.2%, DIV56: 86.9 ± 8.79%) (Fig. 2, A and C). The expression of ACAN, however, is different (Fig. 2, B and C). ACAN staining is present in a small proportion of neurons at DIV21 (19.1 ± 11.9%), and significantly increased by DIV28 with more than doubling to account for 52.5 ± 5.63% of all PNNs, and further increase to 94.3 ± 6.45% at DIV56. This indicates that WFA is a better marker than ACAN for following PNN development.

**Figure 2.**
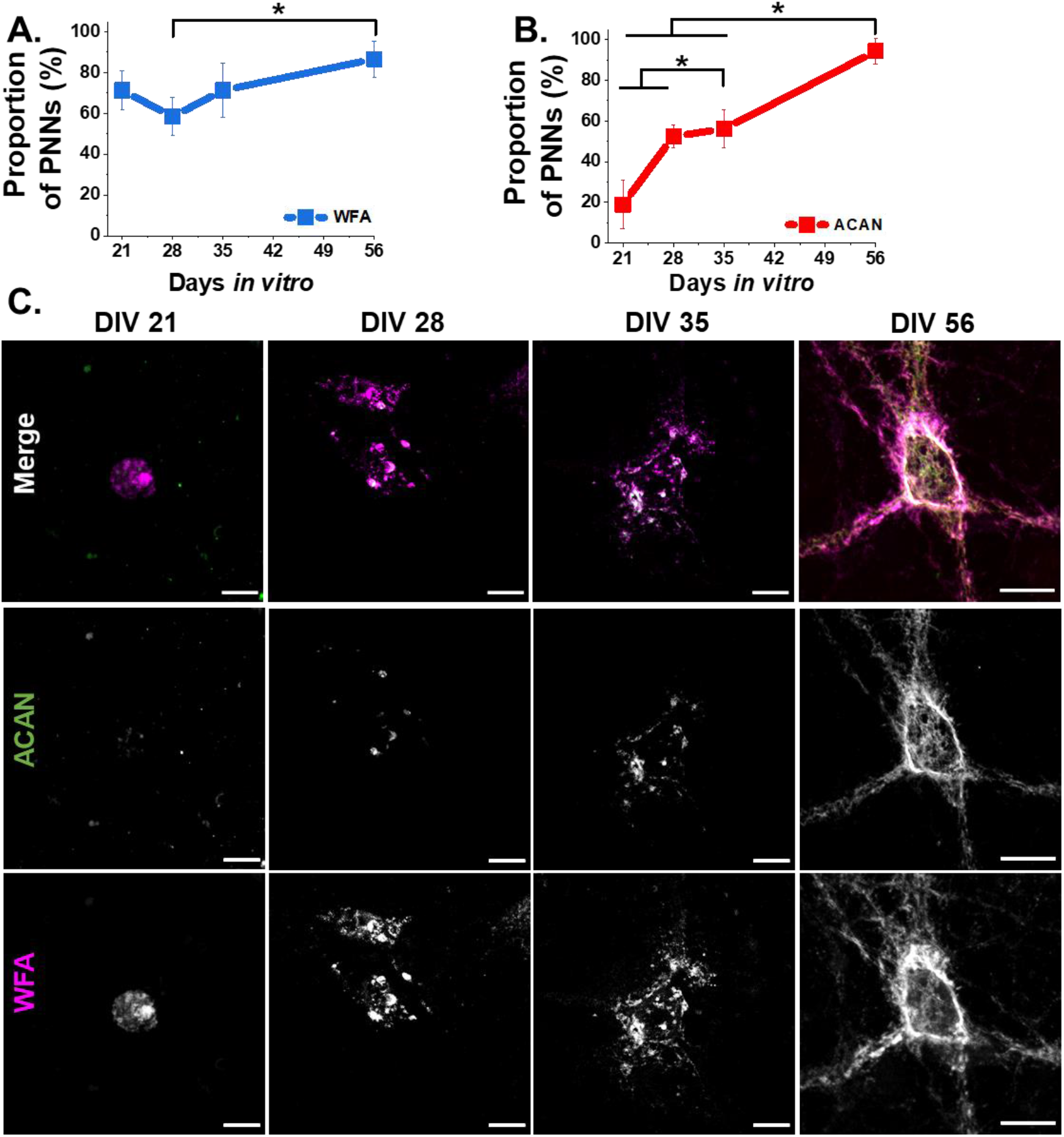
Recruitment and colocalisation of PNN components. **(A)** Percentage of PNNs that are WFA positive during maturation *in vitro*. **(B)** Percentage of PNNs that are ACAN positive during maturation *in vitro.* Data are the average percentages of PNN subtype arising from 6 coverslips from 2 rats. **(C)** Representative images of recruitment of ACAN to punctate, WFA positive PNNs at different time points. Scale bar is 10 μm.

WFA is a lectin that recognises a GAG glycosylation epitope that is enriched in the PNN (Sorg, *et al.*, 2016). The lecticans are the main bearers of GAGs within the PNN and ACAN has been shown to be the main bearer of the WFA-epitope within the cortex (*29*). However, in our culture, we observed this was not the case during PNN development, where ACAN only colocalised with 6.86 ± 1.46% of WFA-positive PNNs at DIV21. ACAN staining began to significantly localise to WFA-positive PNNs at DIV28 and stained increasing proportions of WFA-positive PNNs, with 90.5 ± 5.94% co-location at DIV56. This indicates that another CSPG bears the WFA-epitope at DIV21 rather than ACAN. We were unable to identify the bearer of the WFA epitope. We examined NCAN and VCAN expression but saw no co-localisation.

### 3.3 Identification of a distinct population of PNNs

While the majority of PNNs transitioned from a punctate to reticular morphology during development, we observed a proportion of PNN neurons that did not follow this trend. At DIV21, HAPLN1-positive PNNs already presented with a reticular morphology while those that were WFA-positive were punctate (Fig. 3A). Interestingly, WFA and HAPLN1 staining was completely segregated at this time point, indicating a distinct PNN population (Fig. 3B). The reticular HAPLN1-positive population was also positive for the CSPG neurocan while the punctate WFA-positive population was not (Fig. 3C). This demonstrates the HAPLN1-positive population has a distinct composition as well as morphology.

**Figure 3.**
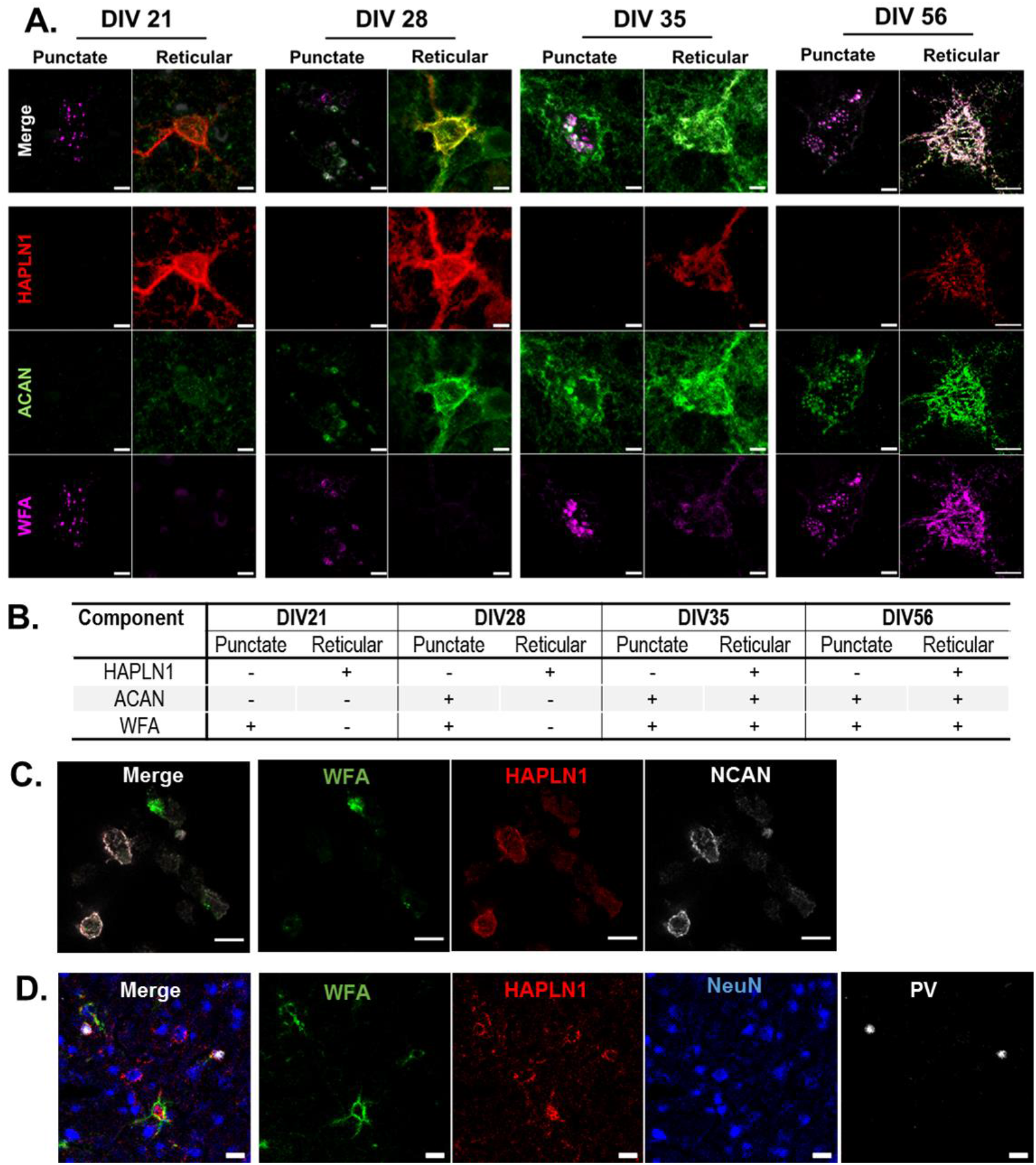
Identification of HAPLN1-positive PNN population. (A) Representative images of PNNs from in vitro culture at DIV28 and (B) summary table. Co-segregation of reticular/HAPLN1 and punctate/WFA neurons during development. Scale bar is 5 μm. (C) HAPLN1-positive PNNs showed co-localisation with NCAN. Scale bar is 10 μm. (D) HAPLN1-positive PNNs are excitatory neurons not PV interneurons. Scale bar is 10 μm.

As mentioned, the localisation of ACAN on PNNs began to increase at DIV28. We observed that ACAN was found on both populations of PNNs (Fig. 3A). ACAN staining on the HAPLN1-positive PNNs bypassed the punctate morphology and presented with a reticular morphology while on WFA-positive PNNs, it presented with the punctate morphology. This would indicate ACAN is localising to an already existent PNN structure which differs in morphology between the PNN populations. WFA staining began to appear on HAPLN1-positive PNNs at DIV35, presenting with a reticular rather than punctate morphology. At no time point in our culture did we observe HAPLN1-positive presenting with a punctate morphology. This could indicate that HAPLN1 is a key protein in determining PNN morphology. This is reminiscent of the *Hapln1* knockout mouse which presents with smaller, diffuse PNNs and persistent plasticity (*8*).

During PNN development, the proportion of HAPLN1-positive PNNs did not change significantly (one-way ANOVA *p*>0.05) and accounted for 31.8 ± 13.7% of total PNNs. This proportion was similar to the proportion of PV-negative PNN neurons in our culture (Fig. 1G). HAPLN1-positive/PV-negative PNN neurons are also observed *in vivo* (Fig. 3D). We therefore hypothesised that there could be overlap between these two populations. In the cortex, the other significant PNN population are excitatory pyramidal neurons. Further work is required to confirm their identity in our culture.

Our results have identified a distinct population of cortical PNNs which present with a different morphology and composition and shown that this population forms on a different neuronal subtype.

### 3.4 Mature in vitro PNNs replicate composition and heterogeneity seen in vivo

At DIV56, the majority of PNNs were reticular in nature, mirroring the mature PNNs seen *in vivo* (Fig. 1). PNN composition *in vivo* is varied with many PNN components identified. We therefore sought to characterise the composition of our mature PNNs *in vitro* to determine whether they replicate that seen *in vivo*. We probed our culture for CSPG, HAPLN1 and tenascin R (TnR) expression as these are key components of PNNs *in vivo*. At DIV56 we identified TnR, HAPLN1 and WFA expression. Out of the CSPGs family lecticans, we observe the presence of ACAN, BCAN and NCAN on the PNNs labelled by WFA (Fig. 4A). We did not observe versican or phosphacan PNN staining on mature PNNs in our culture. The majority of PNNs contained all components stained for (Fig. 4). However, a sizeable proportion of PNNs lacked one or more of the components stained for demonstrating PNN heterogeneity within our culture. HAPLN1, WFA and TnR were found on a similar proportion of PNNs but the CSPG composition of these PNNs differed. HAPLN1 was only found on PNNs that contained all 3 CSPGs stained for while WFA and TnR also identified PNNs that contained fewer CSPGs (Table 1). The lack of clear segregation with HAPLN1 and a particular CSPG argues against a specific interaction for HAPLN1 and CSPGs in the cortex (*42*). WFA showed higher co-localisation with ACAN than any other CSPG (Table 1). ACAN-positive PNNs accounted for 90.5% of the total WFA-positive PNN population indicating that ACAN is the main bearer of the WFA epitope in the mature PNN neurons from our neuronal PNN model. This results agree with previous observation in vivo (14). In contrast, no clear segregation of TnR with a CSPG was observed. This corroborated with the biochemical data that TnR binds to G3 domain in all lecticans. Interestingly, no NCAN only PNNs were observed in our culture while ACAN only and BCAN only PNNs were observed. Together this indicates our neuronal PNN culture model replicates the morphology and heterogeneity in composition observed in vivo.

**Figure 4.**
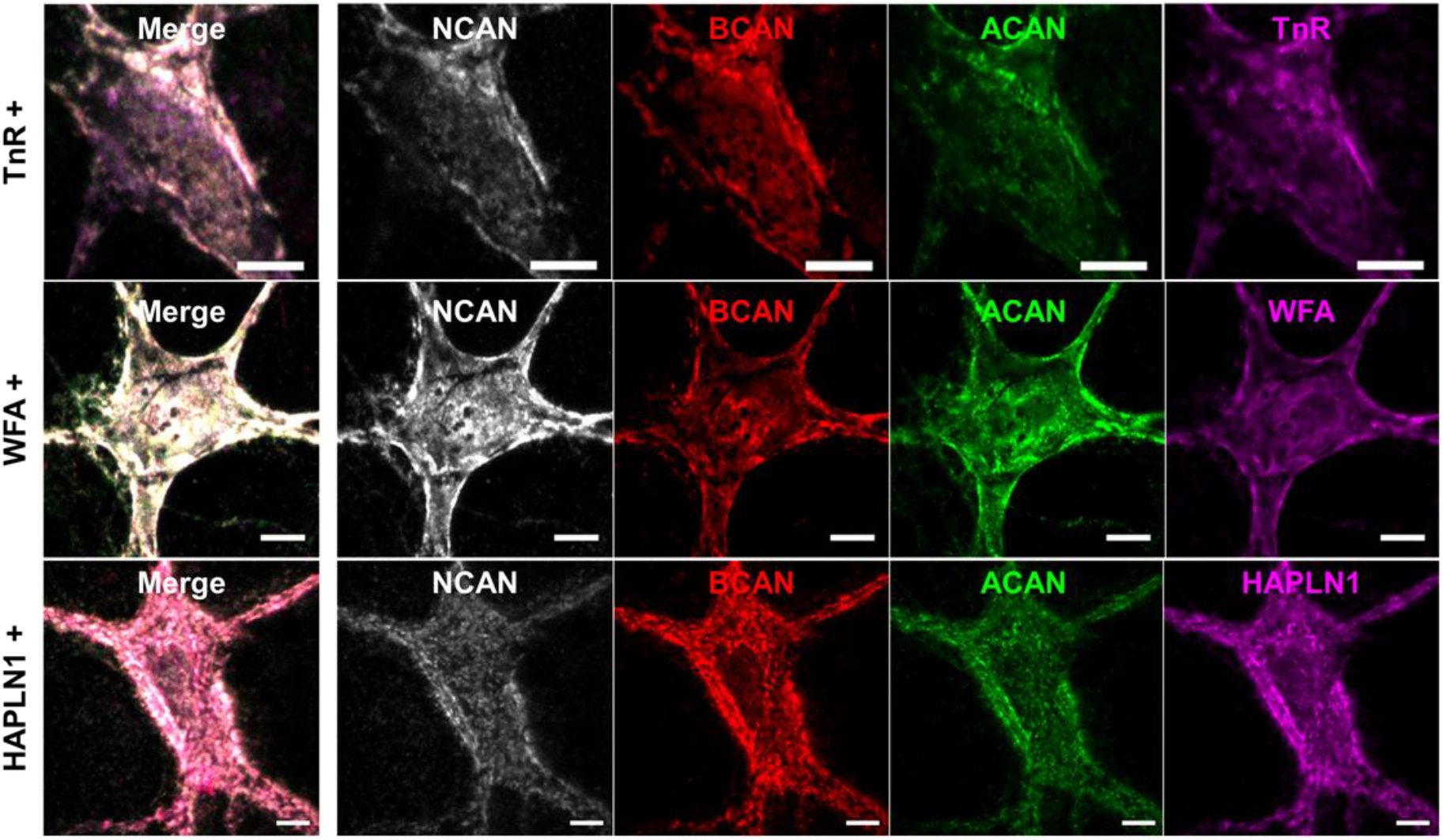
Characterisation of mature PNN cultures. Representative images of DIV56 *in vitro* PNNs containing multiple CSPGs (NCAN, BCAN and ACAN). Triple stained PNNs are also positive for HAPLN1, WFA and TnR. Scale bar is 20 μm.

**Table 1:** Heterogeneity within mature PNNs at DIV56

|  | % of PNNs | ACAN + | NCAN + | BCAN + | % containing 3 CSPGs | % containing 2 CSPGs | % containing 1 CSPG |
| --- | --- | --- | --- | --- | --- | --- | --- |
| HAPLN1 | 79.9 ± 13.8 | 90.3 ± 13.9 | 90.5 ± 10.8 | 90.5 ± 10.8 | 94.4 ± 5.11 | n/a | n/a |
| TnR | 80.7 ± 10.2 | 88.2 ± 19.8 | 75.5 ± 21.5 | 95.8 ± 5.1 | 71.5 ± 22.4 | 16.2 ± 9.72 | 12.5 ± 18.4 |
| WFA | 86.6 ± 8.79 | 90.6 ± 5.94 | 56.6 ± 12.0 | 57.3 ± 17.0 | 49.5 ± 13.5 | 13.9 ± 10.5 | 28.0 ± 16.1 |

**Table 2.** A table of antibodies used.

| Antibody | Species Origin | Dilution | Working concentration (µg/mL) | Company (Product no.) |
| --- | --- | --- | --- | --- |
| **1° antibodies** |  |  |  |  |
| Anti-aggrecan | Rabbit | 1:250 | 2 | Millipore (AB1031) |
| Anti-αSYN | Mouse | 1:250 | 4 | Sigma (S5566) |
| Anti-β<sub>3</sub> Tubulin | Chicken | 1:500 | 0.6 | Abcam (ab41489) |
| Anti-β<sub>3</sub> Tubulin | Mouse | 1:500 | 4 | Abcam (ab6267) |
| Anti-brevican | Sheep | 1:250 | 0.8 | R&D Systems (AF4009) |
| Biotinylated-HABP<sup>1</sup> | - | 1:250 | 1 | AMS Bio (AMS.HKD.B141) |
| Anti-HAPLN1 | Goat | 1:250 | 4 | R&D Systems (AF2608) |
| Anti-HAPLN2 | Rabbit | 1:100 | 0.5 | Novus Biologicals (NBP1-91977) |
| Anti-HAPLN4 | Goat | 1:100 | 10 | Novus Biologicals (AF4085) |
| Anti-NeuN | Guinea-pig | 1:500 | 2 | Synaptic Systems (266004) |
| Anti-neurocan | Mouse | 1:50 | 7.38 | DSHB<sup>2</sup> (1F6) |
| Anti-parvalbumin | Mouse | 1:500 | 2 | Novus Biologicals (NBP2-50038SS) |
| Anti-phosphacan | Mouse | 1:50 | 3.3 | DSHB<sup>2</sup> (3F8) |
| Anti-p129 αSYN | Rabbit | 1:500 | 2 | Abcam (ab51253) |
| Anti-tenascin-R | Goat | 1:250 | 4 | R&D Systems (AF3865) |
| Anti-versican | Mouse | 1:100 | 1.69 | DSHB<sup>2</sup> (12C5) |
| Biotinylated-WFA | - | 1:300 | 6.6 | Sigma (L1516) |
| **2° antibodies** |  |  |  |  |
| Hoechst | - | 1:30,000 | 0.33 | Invitrogen (H3570) |
| CF<sup>TM</sup> 405 M anti-guinea-pig | Donkey | 1:500 | 4 | Sigma (SAB4600468) |
| AF<sup>1</sup> 405 anti-goat | Donkey | 1:500 | 4 | Abcam (Ab175664) |
| Pacific blue streptavidin | - | 1:500 | 4 | Invitrogen (S11222) |
| AF<sup>1</sup> 488 anti-rabbit | Donkey | 1:500 | 4 | Invitrogen (A21206) |
| AF<sup>1</sup> 488 streptavidin | - | 1:500 | 4 | Invitrogen (S32354) |
| AF<sup>1</sup> 568 anti-goat | Donkey | 1:500 | 4 | Invitrogen (A11057) |
| AF<sup>1</sup> 568 anti-mouse | Donkey | 1:500 | 4 | Invitrogen (A10037) |
| AF<sup>1</sup> 568 anti-rabbit | Donkey | 1:500 | 4 | Invitrogen (A10042) |
| AF<sup>1</sup> 568 streptavidin | - | 1:500 | 4 | Invitrogen (S11226) |
| 640/660 Neurotrace<sup>TM</sup> | - | 1:150 | - | Invitrogen (N21483) |
| AF<sup>1</sup> 647 anti-IgY chicken | Goat | 1:500 | 4 | Invitrogen (A21449) |
| AF<sup>1</sup> 647 anti-mouse | Donkey | 1:500 | 4 | Invitrogen (A31571) |

HABP: Hyaluronan binding protein
DSHB: Developmental studies hybridoma bank
AF: Alexa Fluor

## 4.0 Discussion

We have established a PNN culture model that accurately replicates the PNN heterogeneity and complexity observed in the cortex. These neurons mimic the progressive maturation of PNNs in the CNS and thus will serve as a valuable platform that PNN maturation can be studied and manipulated.

The *in vitro* neuronal culture develops mature cortical PNNs at DIV56. WFA reactive PNNs appear around DIV21 and are punctate in nature. Between DIV28-35, the staining coalesces as the culture ages into a mature, reticular morphology. By DIV56 the reticular morphology is the predominant population, accounting for 90% of PNNs, and replicates PNN morphology in the mature CNS. We have shown that WFA is the best marker for tracking PNN development as it stains the majority of PNNs at all-time points. ACAN, the predominant CSPG found in cortical PNNs, appears later in PNN development than WFA and HAPLN1, but localises to the majority of mature PNNs. We have identified two distinct populations of PNNs during development; punctate WFA-positive PNNs (immature) and reticular HAPLN1-positive PNNs (early mature). This distinction disappears as the culture matures and suggests that not all PNNs mature at the same rate. We characterised the composition of mature PNNs at DIV56. We found they contained HAPLN1, TnR and a variety of CSPGs including BCAN, ACAN and NCAN.

Versican and phosphcan were not observed in our neuronal PNN culture model. There are conflicting reports regarding their expression in the PNN *in vitro*. Versican PNN expression was identified in rat hippocampal neurons at DIV21, while we did not observe it our cortical culture at DIV56 (*43*). Phosphacan PNN expression was not found at DIV30 in rat cortical cultures while it has been identified at DIV21 by a different group (*33, 44*). These discrepancies can be explained by temporal and spatial heterogeneity. Versican PNN expression varies between brain region, with higher expression in the hippocampus than the parietal cortex (*31, 39*). Temporal variation in expression has been demonstrated in this culture and elsewhere. Phosphacan PNN expression has been shown early in PNN development (*14, 45*). Another possibility for their absence is that these components are produce by other glial cells in response to the neuronal signals, which would be missed in our astrocyte-conditioned culture medium.

### 4.1 PNN maturation correlates with increasing ACAN localisation

During development, the PNN transitions from a punctate to a reticular morphology (*27, 35*). In our culture, we observe PNN maturation as a punctate morphology gives way to a reticular structure. The punctate form has also been shown in other neuronal cell culture models at similar time points *in vitro* (*22, 32, 33*), but this is the first time the transition to a reticular morphology has been reported. We observed that PNN maturation enters a key point between DIV28-35 as this is when the PNN population switches from the immature state to the mature. ACAN signal increases during this period of *in vitro* maturation, indicating a role in the maturation process. Indeed, conditional knockout mice of ACAN in the CNS show no PNNs and a high level of plasticity in ocular dominance plasticity (*29*). The expression of ACAN mRNA increases during PNN maturation in several CNS regions, including the parietal cortex (*14, 16, 31*).

### 4.2 An early mature PNN population

WFA reactivity started appearing at DIV21 in our culture and was punctate in appearance, accounting for 71.3 ± 4.88% of PNNs. A HAPLN1-positive reticular PNN population was also observed which accounted for the remaining PNNs. They are distinct populations due to the minimal overlap between the staining (HAPLN1+/WFA+: 3.38 ± 2.15%, WFA+/HAPLN1+: 8.02 ± 3.12%) compared to the proportion of WFA-positive PNNs that were ACAN-positive at DIV21. As the culture matured, the co-localisation of ACAN, HAPLN1 and WFA significantly increased. The identity of this population was not resolved during this project but is likely that they are non-PV neurons as the proportion of PNN neurons that were not PV neurons was also around 30%. In the cortex, excitatory pyramidal neurons are the second commonest population of PNN positive neurons (*46–48*). Reticular PNNs have been shown to form on cortical pyramidal neurons *in vivo (49–51)*. Excitatory circuits typically mature before their inhibitory counterparts and help driving their maturation in the cortex (*52*). The earlier maturation of excitatory neurons could therefore explain the early appearance of mature PNNs.

### 4.3 HAPLN1 does not segregate with a specific CSPG

When the HAPLN family were first identified, it was noted that each member showed genetic linkage to a lectican gene (*42*). This led to the hypothesis that each HAPLN protein had a preferential CSPG partner. This hypothesis would other a neat explanation to the apparent redundancy within the HAPLN family but has yet to be proved. Data from *Hapln* knockout animals are conflicted. *Hapln1* knockout resulted in the loss of all CSPGs from cortical PNNs, despite the fact *Hapln4* is also found in cortical PNNs (*8, 19, 29*). The protein expression of the CSPGs was conserved in the knockout, indicating it was the structure the components bound to rather than the components themselves that was lost (*8*). This would indicate there is no specific HAPLN/CSPG pairing. However, in the *Hapln4* knockout animal, BCAN expression was completely lost from cerebellar PNNs, while HAPLN1/ACAN localisation was unaffected. Also, NCAN was only partially affected (*53*). This indicates a preferential pairing between HAPLN1, NCAN and ACAN. This would indicate a specific pairing between HAPLN4/BCAN in the cerebellum but it is in contrast to the visual cortex where *Hapln1* knockout caused ablation of BCAN localisation to the PNNs (*8*). In our cortical culture, we saw no segregation of HAPLN1 with a single CSPG. Instead, it was only found in PNNs with all three CSPGs. This agrees with the *Hapln1* knockout mouse model which also showed no preference. This shows that there is no specific pairing of HAPLN1 with a CSPG in the cortex. However, this does not reconcile with what is seen in cerebellar PNNs which indicates organisational differences between PNNs in different CNS regions. To resolve this the biophysical interactions between HAPLNs and CSPGs should be systematically screened.

### 4.4 TnR localises to mature PNNs in vitro

TnR is a necessary component of PNNs as its removal leads to punctate and reduced PNNs (*25, 54*). This is reminiscent of the immature punctate PNNs seen in our culture and *in vivo*. TnR stained a high proportion of PNNs and seemed to segregate from ncan as no TnR-positive/NCAN only PNNs were identified. This is interesting as an interaction between NCAN and TnR has been demonstrated (*55*).

## 5.0 Conclusion

Here, we provide evidence for a neuronal culture model which accurately modelled the *in vivo* development of PNNs, both in terms of their molecular heterogeneity and morphology. The PNNs in this culture replicate the mature, reticular morphology seen in the mature CNS. We have also shown that the PNNs contain the PNN components seen in the cortex: the CSPGs, ACAN, BCAN and NCAN, HAPLN1 and TnR. The *in vivo* PNN heterogeneity is also replicated. The PNN coalesced from a faint punctate morphology to the mature, reticular morphology seen in the adult CNS. Concomitant with this transition is a rise in ACAN staining. Interestingly, we have found a PNN population that was reticular at an earlier time point, indicating different maturation timeframes between PNNs. This culture model which accurately replicates PNN development and heterogeneity would provide a valuable tool for PNN biology and their roles in diseases.

## 6.0 Funding and Acknowledgements

This work was financially supported by the Royal Society (IEC\R3\203086), Leverhulme Trust (RPG-2018_100), European Union (Operational Programme Research, Development and Education) in the framework of the project ‘Centre of Reconstructive Neuroscience’ (CZ.02.1.01/0.0./0.0/15_003/0000419) to JCFK.

The authors wish to thank the BioImaging Facility at the University of Leeds for their advice for the cell imaging.

## References

(1) Lander, C., Kind, P., Maleski, M., and Hockfield, S. (1997) A family of activity-dependent neuronal cell-surface chondroitin sulfate proteoglycans in cat visual cortex. The Journal of neuroscience : the official journal of the Society for Neuroscience. 17, 1928–1939

(2) Adams, I., Brauer, K., Arélin, C., Härtig, W., Fine, A., Mäder, M., Arendt, T., and Brückner, G. (2001) Perineuronal nets in the rhesus monkey and human basal forebrain including basal ganglia. Neuroscience. 108, 285–298

(3) Cornez, G., Jonckers, E., Ter Haar, S.M., Van der Linden, A., Cornil, C.A., and Balthazart, J. (2018) Timing of perineuronal net development in the zebra finch song control system correlates with developmental song learning. Proceedings. Biological sciences. 285,

(4) Edwards, J.A., Risch, M., and Hoke, K.L. (2021) Dynamics of perineuronal nets over amphibian metamorphosis. The Journal of comparative neurology. 529, 1768–1778

(5) Geissler, M., Gottschling, C., Aguado, A., Rauch, U., Wetzel, C.H., Hatt, H., and Faissner, A. (2013) Primary hippocampal neurons, which lack four crucial extracellular matrix molecules, display abnormalities of synaptic structure and function and severe deficits in perineuronal net formation. The Journal of neuroscience : the official journal of the Society for Neuroscience. 33, 7742–7755

(6) Gottschling, C., Wegrzyn, D., Denecke, B., and Faissner, A. (2019) Elimination of the four extracellular matrix molecules tenascin-C, tenascin-R, brevican and neurocan alters the ratio of excitatory and inhibitory synapses. Sci Rep. 9, 13939

(7) Guimarães, A., Zaremba, S., and Hockfield, S. (1990) Molecular and morphological changes in the cat lateral geniculate nucleus and visual cortex induced by visual deprivation are revealed by monoclonal antibodies Cat-304 and Cat-301. The Journal of neuroscience : the official journal of the Society for Neuroscience. 10, 3014–3024

(8) Carulli, D., Pizzorusso, T., Kwok, J.C., Putignano, E., Poli, A., Forostyak, S., Andrews, M.R., Deepa, S.S., Glant, T.T., and Fawcett, J.W. (2010) Animals lacking link protein have attenuated perineuronal nets and persistent plasticity. Brain. 133, 2331–2347

(9) Pizzorusso, T., Medini, P., Landi, S., Baldini, S., Berardi, N., and Maffei, L. (2006) Structural and functional recovery from early monocular deprivation in adult rats. Proceedings of the National Academy of Sciences of the United States of America. 103, 8517–8522

(10) Siebert, J.R., Stelzner, D.J., and Osterhout, D.J. (2011) Chondroitinase treatment following spinal contusion injury increases migration of oligodendrocyte progenitor cells. Exp Neurol. 231, 19–29

(11) Fawcett, J.W., Oohashi, T., and Pizzorusso, T. (2019) The roles of perineuronal nets and the perinodal extracellular matrix in neuronal function. Nature reviews. Neuroscience. 20, 451–465

(12) Rankin-Gee, E.K., McRae, P.A., Baranov, E., Rogers, S., Wandrey, L., and Porter, B.E. (2015) Perineuronal net degradation in epilepsy. Epilepsia. 56, 1124–1133

(13) Carulli, D., Rhodes, K.E., Brown, D.J., Bonnert, T.P., Pollack, S.J., Oliver, K., Strata, P., and Fawcett, J.W. (2006) Composition of perineuronal nets in the adult rat cerebellum and the cellular origin of their components. The Journal of comparative neurology. 494, 559–577

(14) Galtrey, C.M., Kwok, J.C., Carulli, D., Rhodes, K.E., and Fawcett, J.W. (2008) Distribution and synthesis of extracellular matrix proteoglycans, hyaluronan, link proteins and tenascin-R in the rat spinal cord. Eur J Neurosci. 27, 1373–1390

(15) Giamanco, K.A., and Matthews, R.T. (2012) Deconstructing the perineuronal net: cellular contributions and molecular composition of the neuronal extracellular matrix. Neuroscience. 218, 367–384

(16) Carulli, D., Rhodes, K.E., and Fawcett, J.W. (2007) Upregulation of aggrecan, link protein 1, and hyaluronan synthases during formation of perineuronal nets in the rat cerebellum. The Journal of comparative neurology. 501, 83–94

(17) Kwok, J.C.F., Carulli, D., and Fawcett, J.W. (2010) In vitro modeling of perineuronal nets: hyaluronan synthase and link protein are necessary for their formation and integrity. J Neurochem. 114, 1447–1459

(18) Eill, G.J., Sinha, A., Morawski, M., Viapiano, M.S., and Matthews, R.T. (2020) The protein tyrosine phosphatase RPTPζ/phosphacan is critical for perineuronal net structure. The Journal of biological chemistry. 295, 955–968

(19) Bekku, Y., Su, W.-D., Hirakawa, S., Fässler, R., Ohtsuka, A., Kang, J.S., Sanders, J., Murakami, T., Ninomiya, Y., and Oohashi, T. (2003) Molecular cloning of Bral2, a novel brain-specific link protein, and immunohistochemical colocalization with brevican in perineuronal nets☆. Molecular and Cellular Neuroscience. 24, 148–159

(20) Favuzzi, E., Marques-Smith, A., Deogracias, R., Winterflood, C.M., Sánchez-Aguilera, A., Mantoan, L., Maeso, P., Fernandes, C., Ewers, H., and Rico, B. (2017) Activity-Dependent Gating of Parvalbumin Interneuron Function by the Perineuronal Net Protein Brevican. Neuron. 95, 639–655.e610

(21) Vo, T., Carulli, D., Ehlert, E.M.E., Kwok, J.C.F., Dick, G., Mecollari, V., Moloney, E.B., Neufeld, G., de Winter, F., Fawcett, J.W., and Verhaagen, J. (2013) The chemorepulsive axon guidance protein semaphorin3A is a constituent of perineuronal nets in the adult rodent brain. Molecular and Cellular Neuroscience. 56, 186–200

(22) Van’t Spijker, H.M., Rowlands, D., Rossier, J., Haenzi, B., Fawcett, J.W., and Kwok, J.C.F. (2019) Neuronal Pentraxin 2 Binds PNNs and Enhances PNN Formation. Neural plasticity. 2019, 6804575

(23) Beurdeley, M., Spatazza, J., Lee, H.H., Sugiyama, S., Bernard, C., Di Nardo, A.A., Hensch, T.K., and Prochiantz, A. (2012) Otx2 binding to perineuronal nets persistently regulates plasticity in the mature visual cortex. The Journal of neuroscience : the official journal of the Society for Neuroscience. 32, 9429–9437

(24) Köppe, G., Brückner, G., Brauer, K., Härtig, W., and Bigl, V. (1997) Developmental patterns of proteoglycan-containing extracellular matrix in perineuronal nets and neuropil of the postnatal rat brain. Cell and tissue research. 288, 33–41

(25) Brückner, G., Grosche, J., Schmidt, S., Härtig, W., Margolis, R.U., Delpech, B., Seidenbecher, C.I., Czaniera, R., and Schachner, M. (2000) Postnatal development of perineuronal nets in wild-type mice and in a mutant deficient in tenascin-R. The Journal of comparative neurology. 428, 616–629

(26) Mauney, S.A., Athanas, K.M., Pantazopoulos, H., Shaskan, N., Passeri, E., Berretta, S., and Woo, T.U. (2013) Developmental pattern of perineuronal nets in the human prefrontal cortex and their deficit in schizophrenia. Biol Psychiatry. 74, 427–435

(27) Ueno, H., Suemitsu, S., Okamoto, M., Matsumoto, Y., and Ishihara, T. (2017) Sensory experience-dependent formation of perineuronal nets and expression of Cat-315 immunoreactive components in the mouse somatosensory cortex. Neuroscience. 355, 161–174

(28) Sigal, Y.M., Bae, H., Bogart, L.J., Hensch, T.K., and Zhuang, X. (2019) Structural maturation of cortical perineuronal nets and their perforating synapses revealed by superresolution imaging. Proceedings of the National Academy of Sciences of the United States of America. 116, 7071–7076

(29) Rowlands, D., Lensjø, K.K., Dinh, T., Yang, S., Andrews, M.R., Hafting, T., Fyhn, M., Fawcett, J.W., and Dick, G. (2018) Aggrecan Directs Extracellular Matrix-Mediated Neuronal Plasticity. The Journal of Neuroscience. 38, 10102–10113

(30) Hirakawa, S., Oohashi, T., Su, W.D., Yoshioka, H., Murakami, T., Arata, J., and Ninomiya, Y. (2000) The brain link protein-1 (BRAL1): cDNA cloning, genomic structure, and characterization as a novel link protein expressed in adult brain. Biochem Biophys Res Commun. 276, 982–989

(31) Gao, R., Wang, M., Lin, J., Hu, L., Li, Z., Chen, C., and Yuan, L. (2018) Spatiotemporal expression patterns of chondroitin sulfate proteoglycan mRNAs in the developing rat brain. Neuroreport. 29, 517–523

(32) Dityatev, A., Bruckner, G., Dityateva, G., Grosche, J., Kleene, R., and Schachner, M. (2007) Activity-dependent formation and functions of chondroitin sulfate-rich extracellular matrix of perineuronal nets. Dev Neurobiol. 67, 570–588

(33) Miyata, S., Nishimura, Y., Hayashi, N., and Oohira, A. (2005) Construction of perineuronal net-like structure by cortical neurons in culture. Neuroscience. 136, 95–104

(34) Dino, M.R., Harroch, S., Hockfield, S., and Matthews, R.T. (2006) Monoclonal antibody Cat-315 detects a glycoform of receptor protein tyrosine phosphatase beta/phosphacan early in CNS development that localizes to extrasynaptic sites prior to synapse formation. Neuroscience. 142, 1055–1069

(35) Richter, R.P., Baranova, N.S., Day, A.J., and Kwok, J.C.F. (2018) Glycosaminoglycans in extracellular matrix organisation: are concepts from soft matter physics key to understanding the formation of perineuronal nets? Current Opinion in Structural Biology. 50, 65–74

(36) Bardy, C., van den Hurk, M., Eames, T., Marchand, C., Hernandez, R.V., Kellogg, M., Gorris, M., Galet, B., Palomares, V., Brown, J., Bang, A.G., Mertens, J., Böhnke, L., Boyer, L., Simon, S., and Gage, F.H. (2015) Neuronal medium that supports basic synaptic functions and activity of human neurons in vitro. Proceedings of the National Academy of Sciences of the United States of America. 112, E2725–2734

(37) Brückner, G., Hausen, D., Härtig, W., Drlicek, M., Arendt, T., and Brauer, K. (1999) Cortical areas abundant in extracellular matrix chondroitin sulphate proteoglycans are less affected by cytoskeletal changes in Alzheimer’s disease. Neuroscience. 92, 791–805

(38) Mueller, A.L., Davis, A., Sovich, S., Carlson, S.S., and Robinson, F.R. (2016) Distribution of N-Acetylgalactosamine-Positive Perineuronal Nets in the Macaque Brain: Anatomy and Implications. Neural plasticity. 2016, 6021428

(39) Ueno, H., Suemitsu, S., Okamoto, M., Matsumoto, Y., and Ishihara, T. (2017) Parvalbumin neurons and perineuronal nets in the mouse prefrontal cortex. Neuroscience. 343, 115–127

(40) Irvine, S.F., and Kwok, J.C.F. (2018) Perineuronal Nets in Spinal Motoneurones: Chondroitin Sulphate Proteoglycan around Alpha Motoneurones. International Journal of Molecular Sciences. 19, 1172

(41) Morawski, M., Brückner, G., Jäger, C., Seeger, G., Matthews, R.T., and Arendt, T. (2012) Involvement of perineuronal and perisynaptic extracellular matrix in Alzheimer’s disease neuropathology. Brain pathology (Zurich, Switzerland). 22, 547–561

(42) Spicer, A.P., Joo, A., and Bowling, R.A., Jr. (2003) A hyaluronan binding link protein gene family whose members are physically linked adjacent to chondroitin sulfate proteoglycan core protein genes: the missing links. The Journal of biological chemistry. 278, 21083–21091

(43) John, N., Krugel, H., Frischknecht, R., Smalla, K.H., Schultz, C., Kreutz, M.R., Gundelfinger, E.D., and Seidenbecher, C.I. (2006) Brevican-containing perineuronal nets of extracellular matrix in dissociated hippocampal primary cultures. Molecular and cellular neurosciences. 31, 774–784

(44) Giamanco, K.A., Morawski, M., and Matthews, R.T. (2010) Perineuronal net formation and structure in aggrecan knockout mice. Neuroscience. 170, 1314–1327

(45) Hayashi, N., Oohira, A., and Miyata, S. (2005) Synaptic localization of receptor-type protein tyrosine phosphatase zeta/betain the cerebral and hippocampal neurons of adult rats. Brain Res. 1050, 163–169

(46) Seeger, G., Brauer, K., Härtig, W., and Brückner, G. (1994) Mapping of perineuronal nets in the rat brain stained by colloidal iron hydroxide histochemistry and lectin cytochemistry. Neuroscience. 58, 371–388

(47) Pizzorusso, T., Medini, P., Berardi, N., Chierzi, S., Fawcett, J.W., and Maffei, L. (2002) Reactivation of ocular dominance plasticity in the adult visual cortex. Science. 298, 1248–1251

(48) Brückner, G., Kacza, J., and Grosche, J. (2004) Perineuronal Nets Characterized by Vital Labelling, Confocal and Electron Microscopy in Organotypic Slice Cultures of Rat Parietal Cortex and Hippocampus. Journal of Molecular Histology. 35, 115–122

(49) Hausen, D., Brückner, G., Drlicek, M., Härtig, W., Brauer, K., and Bigl, V. (1996) Pyramidal cells ensheathed by perineuronal nets in human motor and somatosensory cortex. Neuroreport. 7, 1725–1729

(50) Wegner, F., Härtig, W., Bringmann, A., Grosche, J., Wohlfarth, K., Zuschratter, W., and Brückner, G. (2003) Diffuse perineuronal nets and modified pyramidal cells immunoreactive for glutamate and the GABAA receptor α1 subunit form a unique entity in rat cerebral cortex. Experimental Neurology. 184, 705–714

(51) Alpar, A., Gartner, U., Hartig, W., and Bruckner, G. (2006) Distribution of pyramidal cells associated with perineuronal nets in the neocortex of rat. Brain Res. 1120, 13–22

(52) Lodato, S., Rouaux, C., Quast, K.B., Jantrachotechatchawan, C., Studer, M., Hensch, T.K., and Arlotta, P. (2011) Excitatory projection neuron subtypes control the distribution of local inhibitory interneurons in the cerebral cortex. Neuron. 69, 763–779

(53) Bekku, Y., Saito, M., Moser, M., Fuchigami, M., Maehara, A., Nakayama, M., Kusachi, S., Ninomiya, Y., and Oohashi, T. (2012) Bral2 is indispensable for the proper localization of brevican and the structural integrity of the perineuronal net in the brainstem and cerebellum. The Journal of comparative neurology. 520, 1721–1736

(54) Morawski, M., Dityatev, A., Hartlage-Rübsamen, M., Blosa, M., Holzer, M., Flach, K., Pavlica, S., Dityateva, G., Grosche, J., Brückner, G., and Schachner, M. (2014) Tenascin-R promotes assembly of the extracellular matrix of perineuronal nets via clustering of aggrecan. Philosophical Transactions of the Royal Society of London B: Biological Sciences. 369, 20140046

(55) Aspberg, A., Miura, R., Bourdoulous, S., Shimonaka, M., Heinegård, D., Schachner, M., Ruoslahti, E., and Yamaguchi, Y. (1997) The C-type lectin domains of lecticans, a family of aggregating chondroitin sulfate proteoglycans, bind tenascin -R by protein– protein interactions independent of carbohydrate moiety. Proceedings of the National Academy of Sciences. 94, 10116–10121

